# The highly expressed ERV1 forms virus-like particles for regulating early embryonic development

**DOI:** 10.1101/2022.06.21.496818

**Authors:** Wenjing Li, Shujuan Liu, Jianglin Zhao, Ruizhi Deng, Yayi Liu, Huijia Li, Hongwei Ma, Yanzhi Chen, Jingcheng Zhang, Yongsheng Wang, Jianmin Su, Fusheng Quan, Xu liu, Yan Luo, Yong Zhang, Jun Liu

**Affiliations:** College of Veterinary Medicine, Northwest A&F University, Key Laboratory of Animal Biotechnology of the Ministry of Agriculture, Yangling 712100, Shaanxi, China; Department of Animal Engineering, Yangling Vocational and Technical College, Yangling 712100, China; Reproductive hospital, Jiangxi University of Traditional Chinese Medicine, Nanchang 330004, Jiangxi, China

**Keywords:** transposable elements, endogenous retrovirus, virus-like particles, trophoblast cell differentiation, transcriptome sequences, embryonic development

## Abstract

In mammals, the transcription of transposable elements (TEs) is important for maintaining early embryonic development. Here, we systematically analyzed the expression characteristics of TE-derived transcripts in early embryos by constructing a database of TEs and transcriptome data from goats and using it to study the function of endogenous retroviruses (ERVs) in regulating early embryo development. We found that ERV1 made up the highest proportion of TE sequences and exhibited a stage-specific expression pattern during early embryonic development. Among ERV elements, ERV1 had the potential to encode the Gag protein domain to form virus-like particles (VLPs) in early goat embryos. Knockdown of ERV1_1_574 significantly reduced the embryo development rate and the number of trophoblast cells (*P*< 0.05). Transcriptome sequencing analysis of morula embryos showed that ERV1_1_574 mainly regulated the expression of genes related to embryo compaction and trophoblast cell differentiation, such as CX43 and CDX2. In summary, we found that ERV1 expression was essential for early embryonic development in goats through regulation of trophoblast cell differentiation.

## Introduction

TEs are mobile genomic DNA sequences, which are ubiquitous to all organisms and are an important part of the host genome. In humans (44%), mice (40%), cattle (46%) and other organisms, TEs account for almost half of the host genome (Lander *et al*, 2001; Waterston *et al*, 2002; Adelson *et al*, 2009), and increasing evidence indicates that TEs have important effects on the structure, function and evolutionary dynamics of the genome (Xu & Wang, 2007; Rho & Tang, 2009; Senft & Macfarlan, 2021). Recent studies have reported that a large number of transcripts of TEs have been discovered at specific stages of early embryonic development, but their biological functions and regulatory mechanisms have not been fully determined (Gifford *et al*, 2013; Fu *et al*, 2019).

Whole-genome sequencing revealed high variability in the number, type, and sequence of TEs despite the high sequence homology of functional genes in mammals (Platt *et al*, 2018). At present, the TE sequences of a variety of animals can be retrieved from the Repbase database, but the sequences of TEs in the goat genome have not been included or reported yet. Determining the functions and regulatory mechanisms of TEs in early embryonic development of domestic animals is the key to revealing their characteristics in early embryo development. Studies have demonstrated that LINE1 (Beraldi *et al*, 2006), MuERV-L (Kigami *et al*, 2003), LincGET (Wang *et al*, 2018) and HPAT5 (Durruthy-Durruthy *et al*, 2016), among others, are essential regulatory elements for maintaining early embryonic development.

ERV elements constitute a large part of the TEs in eukaryotic genomes (Makalowski *et al*, 2012), accounting for about ten percent of the mammalian genome (Rebollo *et al*, 2012), and many studies showed that ERVs were important for regulating mammalian gene expression (Friedli & Trono, 2015; Thompson *et al*, 2016). Genes of the mouse endogenous retrovirus-like (MERVL) virus initiate a large number of specific transcripts at the two-cell embryo stage (Kigami *et al*, 2003; Macfarlan *et al*, 2012). For example, the ERV-related lncRNA, LincGET, is indispensable for cell division in mouse embryos (Wang *et al*, 2016). In some ERVs, the Gag sequence has the capacity to encode proteins, and the Gag protein forms VLPs that can infect and transmit information between cells, thus performing a regulatory and communication function. In two-cell stage mouse embryos, some MERVL virus sequences can also encode Gag protein (Macfarlan *et al*, 2012). During Drosophila oogenesis, ERV elements in trophoblasts express VLPs, which can infect oocytes and may regulate oogenesis (Wang *et al*, 2018). Many mammalian ERV element transcripts have been discovered in early embryos, but the mechanism of their regulation of embryo development needs further clarification.

In our previous study, we found that ERV-derived lncRNAs affect the development rate of early goat embryo blastocysts by regulating the expression of the target gene, CHD1L (Deng *et al*, 2019). In this study, a database of TEs in the goat genome was established, and the transcriptome of early goat embryos at various stages was generated by RNA-seq. We systematically analyzed the transcripts of TEs in early goat embryos, and focused on the expression and function of ERV1. We found that ERV1_1_574 was highly expressed in goats during early embryonic development and could encode Gag protein and form virus-like particles. Moreover, ERV1_1_574 could affect embryo development by regulating embryo densification.

## Results

### ERVs are the largest superfamily of TEs in the goat genome

We obtained the sequences of all TEs in the goat genome based on de novo prediction, structure-based prediction and homology-based prediction strategies **(Fig. S1)**. We classified and annotated the TEs to build a comprehensive, user-friendly, web-based database (http://genedenovoweb.ticp.net:81/goatTEdb/) **(Fig.S2)**. A total of 495,065 TEs from 21 super-families and 926 families were included in this database **(Table 1)**. The coverage and classification of TEs in the goat genome were determined by RepeatMasker. We found that the percentage of the goat genome consisting of TEs was similar to that of cattle at 46.33%. LINEs accounted for 10.31% of the genome, which was lower than for cattle (23.29%), humans (20.40%) and mice (19.59%). SINEs made up 2.86% of the goat genome, compared to 17.66% for cattle, 13.11% for humans, and 7.34% for mice. ERVs constituted 28.45% of the goat genome, which was significantly higher than that of cattle (3.20%), humans (8.56%) and mice (9.84%). ERV1 had the highest copy number (12,366 members) among all subclasses in the goat genome **(Table 2)**. The goat TEs were evenly distributed in every chromosome except chromosome 10 **(Fig. S3A)**. Using the sequences of the intact elements to construct a maximum likelihood tree, we identified one BovB, 91 L1, and 41 ERV1 active elements **(Fig. S3B)**. The high proportion of ERVs revealed by TE analysis was a significant feature of the goat genome, in which the ERV1 copy number was particularly high and contained a large proportion of active elements.

**Table 1.**
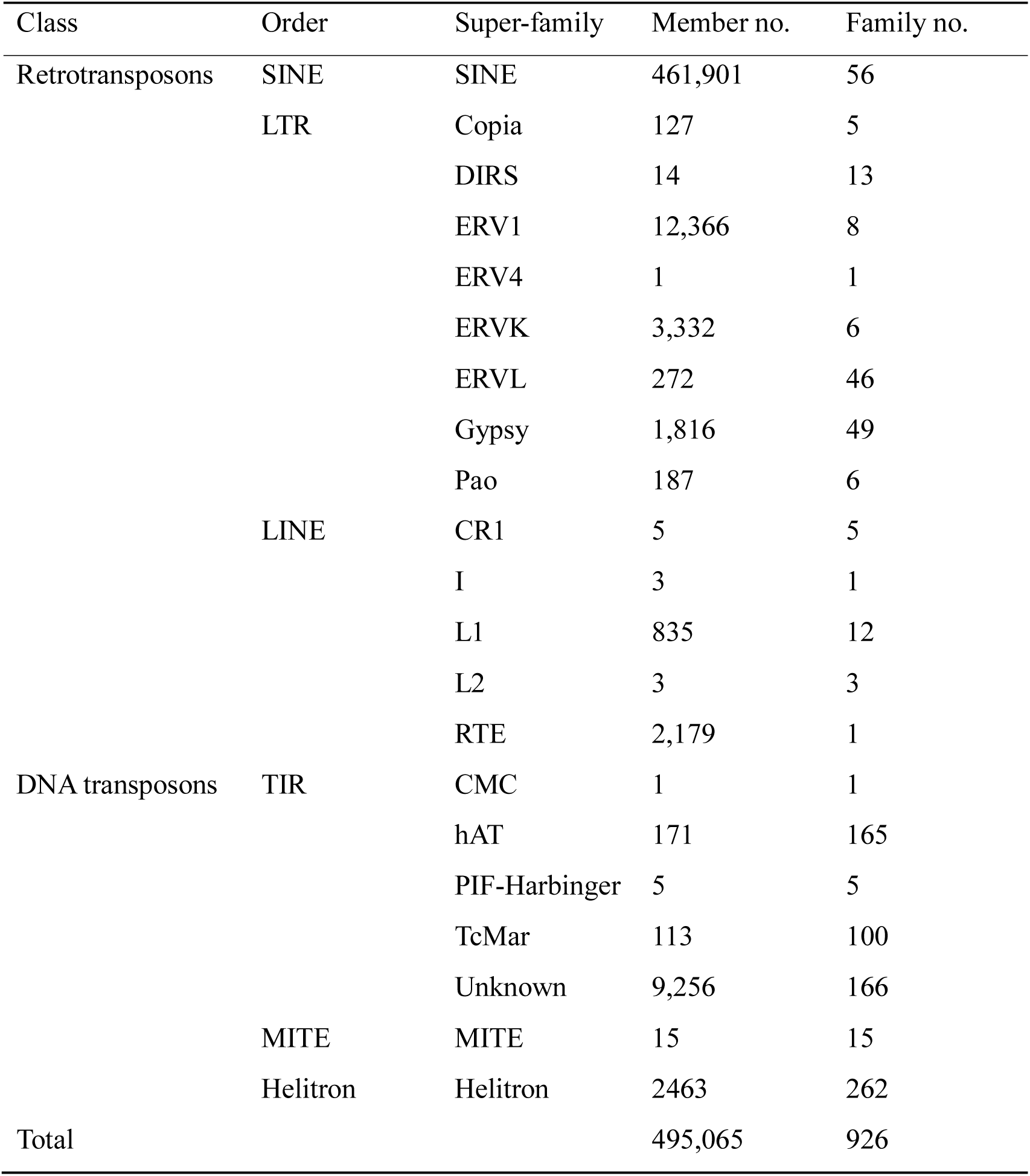
The TEs identified in the goat genome.

**Table 2.**
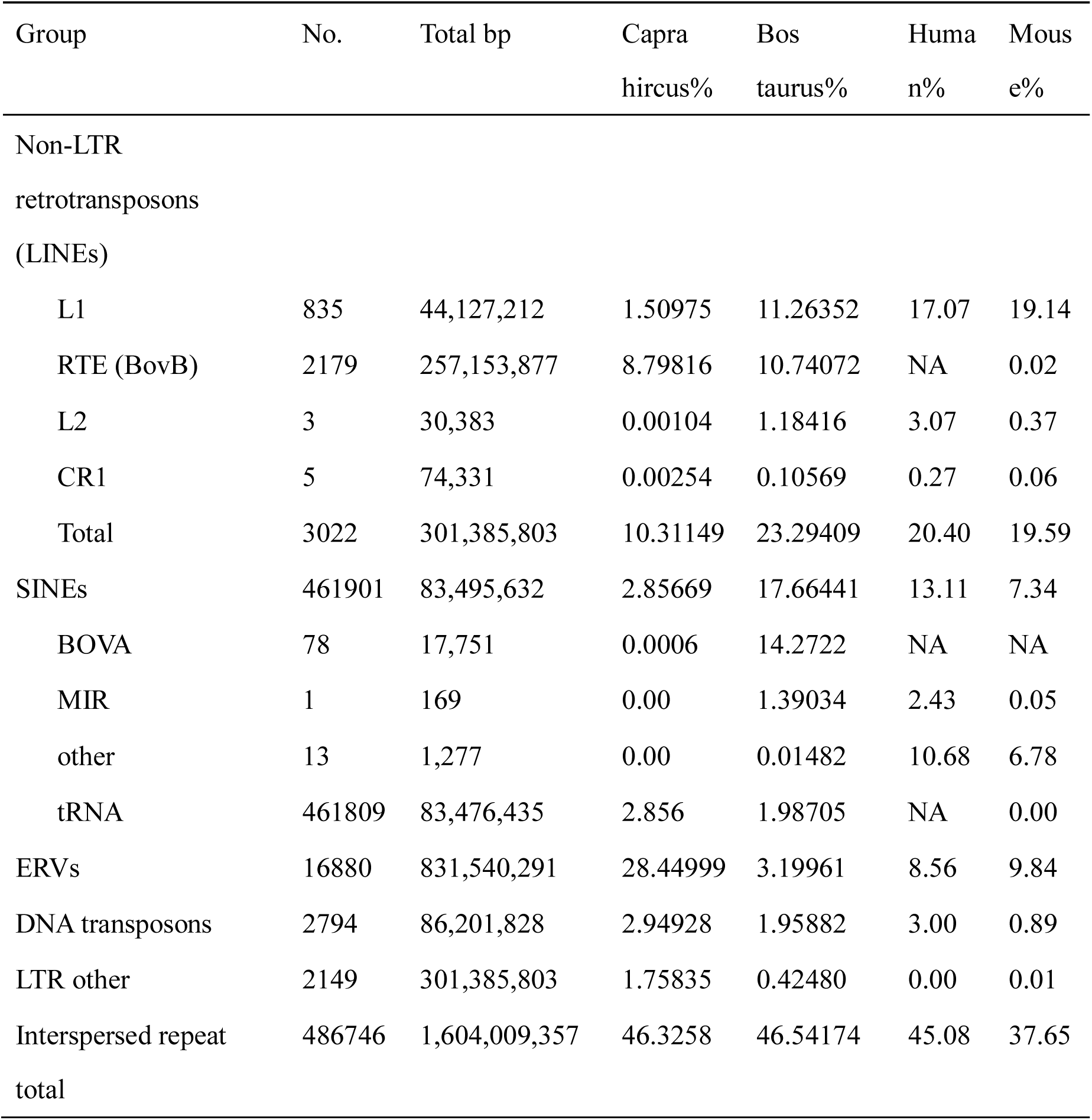
Proportion of TEs in goat genome compared with cattle, human and mouse.

| Group | No. | Total bp | Capra<br>hircus% | Bos<br>taurus% | Huma<br>n% | Mous<br>e% |
| --- | --- | --- | --- | --- | --- | --- |
| Non-LTR<br>retrotransposons<br>(LINEs) |  |  |  |  |  |  |
| L1 | 835 | 44,127,212 | 1.50975 | 11.26352 | 17.07 | 19.14 |
| RTE (BovB) | 2179 | 257,153,877 | 8.79816 | 10.74072 | NA | 0.02 |
| L2 | 3 | 30,383 | 0.00104 | 1.18416 | 3.07 | 0.37 |
| CR1 | 5 | 74,331 | 0.00254 | 0.10569 | 0.27 | 0.06 |
| Total | 3022 | 301,385,803 | 10.31149 | 23.29409 | 20.40 | 19.59 |
| SINEs | 461901 | 83,495,632 | 2.85669 | 17.66441 | 13.11 | 7.34 |
| BOVA | 78 | 17,751 | 0.0006 | 14.2722 | NA | NA |
| MIR | 1 | 169 | 0.00 | 1.39034 | 2.43 | 0.05 |
| other | 13 | 1,277 | 0.00 | 0.01482 | 10.68 | 6.78 |
| tRNA | 461809 | 83,476,435 | 2.856 | 1.98705 | NA | 0.00 |
| ERVs | 16880 | 831,540,291 | 28.44999 | 3.19961 | 8.56 | 9.84 |
| DNA transposons | 2794 | 86,201,828 | 2.94928 | 1.95882 | 3.00 | 0.89 |
| LTR other | 2149 | 301,385,803 | 1.75835 | 0.42480 | 0.00 | 0.01 |
| Interspersed repeat<br>total | 486746 | 1,604,009,357 | 46.3258 | 46.54174 | 45.08 | 37.65 |

### Transcriptome analysis of early goat embryos using RNA-seq

We collected in vitro fertilized (IVF) early goat embryos at various stages **(Fig. 1A)**, and used RNA-seq technology to obtain the transcriptomes. We aligned the reads to the goat genome and assembled the transcripts to analyze mRNA and potential lncRNA expression in early goat embryos **(Fig. S1, Table 3)**. Principal component analysis (PCA) showed that the different developmental stages of embryos could be separated into distinct groups based on gene counts **(Fig. S4)**. The total expression levels of mRNAs and lncRNAs in 8-cell stage embryos were increased during early embryonic development **(Fig. 1B)**. Large-scale transcriptional activation occurred at the 8-cell stage, as expected for embryonic genome activation (EGA) of early goat embryos. The heatmap of dynamic gene expression showed that differentially expressed genes (DEGs) in 8-cell embryos and morula embryos were different from the zygote, 2-cell and 4-cell stage embryos **(Fig. 1C)**. To further explore the dynamic expression patterns of DEGs, we divided all developmental stages into three modules **(Fig. 1D)**. The first module included up-regulated genes in zygotic, 2-cell and 4-cell stages. These were defined as maternal genes and a total of 41 were identified, including 15 lncRNAs and 26 mRNAs, which were enriched in cell division, cell cycle, mitosis, and mitotic cell cycle G/2M transitions. The second module contained highly expressed genes in 8-cell stage embryos. There were 51 genes in total, including 25 lncRNAs and 26 mRNAs. Notably, the typical ZSCAN4 zygotically-activated genes in mice and humans were in this module. GO enrichment indicated that genes in the second module were related to protein binding, DNA binding regulation and GTPase activation. The third module included highly co-expressed genes of 8-cell stage and morula embryos, and 28 genes were screened, including 12 lncRNAs and 16 mRNAs. Genes were mainly enriched for transcription factor activity, transcriptional regulation and transcription-related entries. The second and third modules were collectively defined as zygotic genes, mainly related to transcriptional and regulatory pathways.

**Figure 1.**
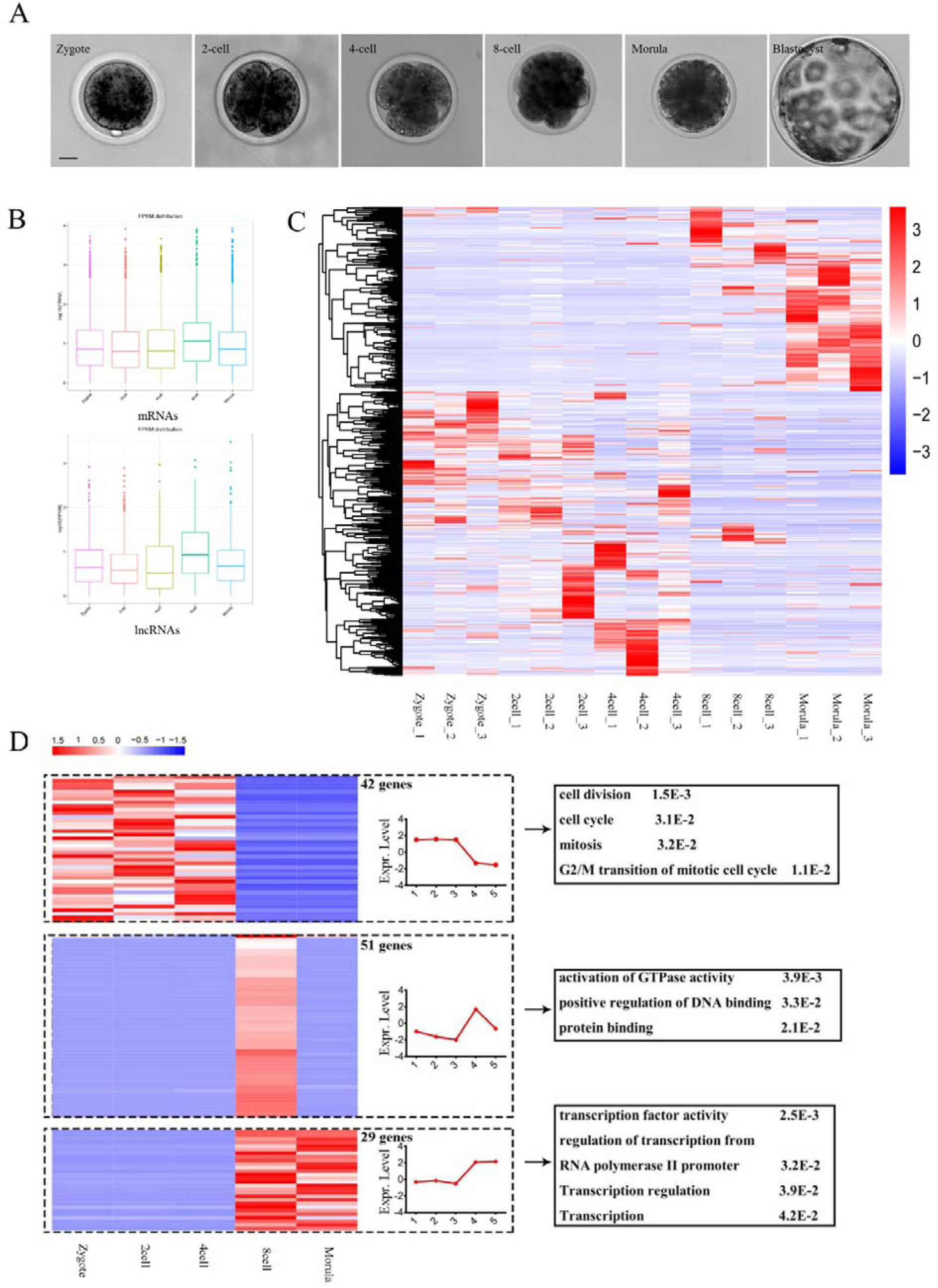
Transcriptome dynamics in early goat embryos. **(A)** Microscopy images of goat pre-implantation embryos at zygote, two-, four-, and eight-cell stages, morulae and blastocysts. Scale bar=20 μm. **(B)** The mRNA and lncRNA expression patterns during goat preimplantation development. **(C)** Hierarchical clustering analysis shows stage-specific (|fold change| >2, *P*< 0.05) expression of genes in goat preimplantation embryo samples. Differentially-expressed genes (DEGs) were appropriately aggregated into different clusters. Red: high expression. Blue: low expression. **(D)** Cluster analysis of DEGs. Clusters of DEGs by normalized FPKM reflect the total transcript content per sample during preimplantation development. All of the genes shown were differentially expressed between two consecutive stages (|fold change| >2, *P*< 0.05). The DEGs can be classified as follows: (1) 42 maternal genes with two to three clusters activated in the 8-cell stage defined as zygotic genes; (2) 51 zygotic genes, and (3) 29 zygotic genes. The expression level is shown in the middle panel and GO enrichment is shown in the right panel.

**Table 3.** Number of mRNAs and lncRNAs in goat transcriptome.

| Group | Known | New | All | Known | New | All |
| --- | --- | --- | --- | --- | --- | --- |
| name | mRNA | mRNA | mRNA | lncRNA | lncRNA | lncRNA |
|  | Num | Num | Num | Num | Num | Num |
| Zygote | 12,407 | 6,895 | 19,302 | 247 | 1,862 | 2,109 |
| 2cell | 13,701 | 7,380 | 21,081 | 291 | 2,239 | 2,530 |
| 4cell | 15,444 | 7,912 | 23,356 | 355 | 2,596 | 2,951 |
| 8cell | 10,934 | 5,775 | 16,709 | 234 | 1,103 | 1,337 |
| Morula | 13,246 | 6,366 | 19,612 | 357 | 1,094 | 1,451 |

### ERV1 exhibits a highly expressed, stage-specific pattern in early goat embryos

ERVs were the most abundant class of TE in the goat genome in comparison with bovine, human and mouse genomes **(Table 2, Fig. 2A)**. The expression pattern of TE-derived transcripts in the transcriptome data from each developmental stage of early goat embryos was analyzed **(Fig. S1)**. We found that ERV1s accounted for the highest proportion in the transcriptome at all stages (the top six TEs expressed) **(Fig. 2B)**. Moreover, the number of specifically expressed ERV1s significantly increased in the 8-cell stage embryos when focusing on the ERV1, RTE, and L1 specifically expressed in embryos at different developmental stages (*P*< 0.01) **(Fig. 2C)**. We aligned transcriptome sequences with TE databases to obtain dynamic expression levels of transcripts associated with transposon elements. PCA **(Fig. S5A**) and dynamic expression analysis of all screened stage-specific elements showed that the TEs (ERV1, L1 and RTE) were stage-specifically expressed during development of early goat embryos **(Fig. 2D and S5A)**. We selected the representative maternal gene expression element ERV1_1_2471 and the zygotic gene expression element ERV1_1_12704 for IGV visualization **(Fig. S5B)**, and the sequencing data were verified by RT-qPCR **(Fig. S5C)**. These findings suggested that ERV1s were highly expressed in early goat embryos and exhibited stage-specific expression patterns.

**Figure 2.**
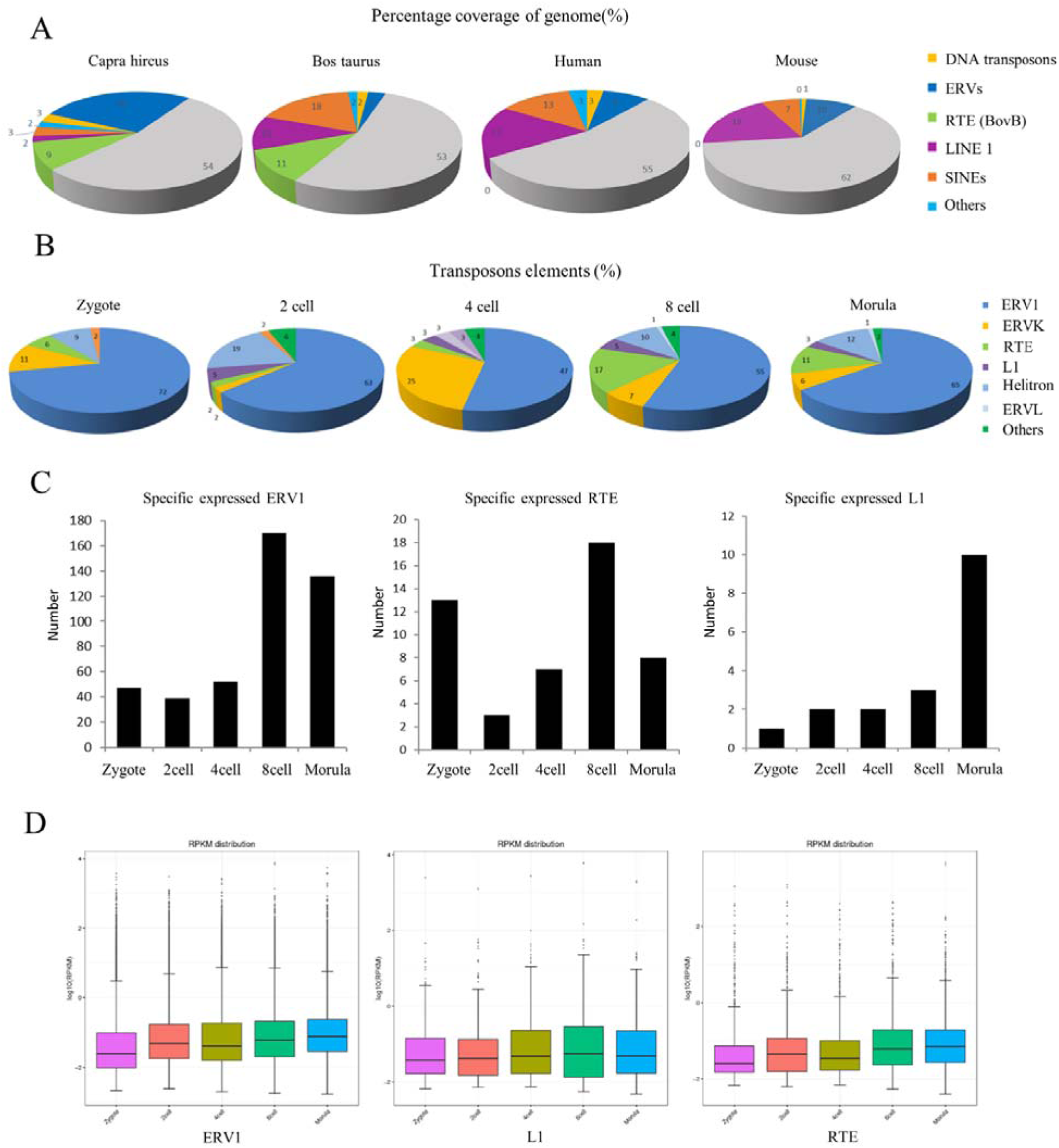
ERV1 is a specific element in goat TE-derived transcripts. **(A)** The proportion of TEs in genomes of different species. **(B)** Relative abundance of TEs at different developmental stages. Relative abundance of transcripts were derived from the first six classes of repetitive elements at the indicated stages. **(C)** The number of ERV1s, RTEs and L1s specifically expressed at different stages compared to other stages. **(D)** Dynamic expression of ERV1, L1 and RTE elements at different stages.

### ERV1 has the potential to form virus-like particles (VLPs)

We performed structural predictions on all ERV1s with complete ERV sequences (domains > 2), and found structures typical of overlapping Gag, Pro and Pol genes in the RNA coding region of ERV1 sequences **(Fig. S6)**. We found that the gag protein structures of ERV1_1_574 and ERV1_1_1613 had the potential to encode a virus-like capsid Gag protein domain P10, P24 and P30 that may generate VLPs **(Fig. 3A)**. The expression level of candidate ERV1 in early embryos was verified by RT-qPCR, and ERV1_1_574 and ERV1_1_1613 were highly expressed at the eight-cell stage **(Fig. 3B)**.

**Figure 3.**
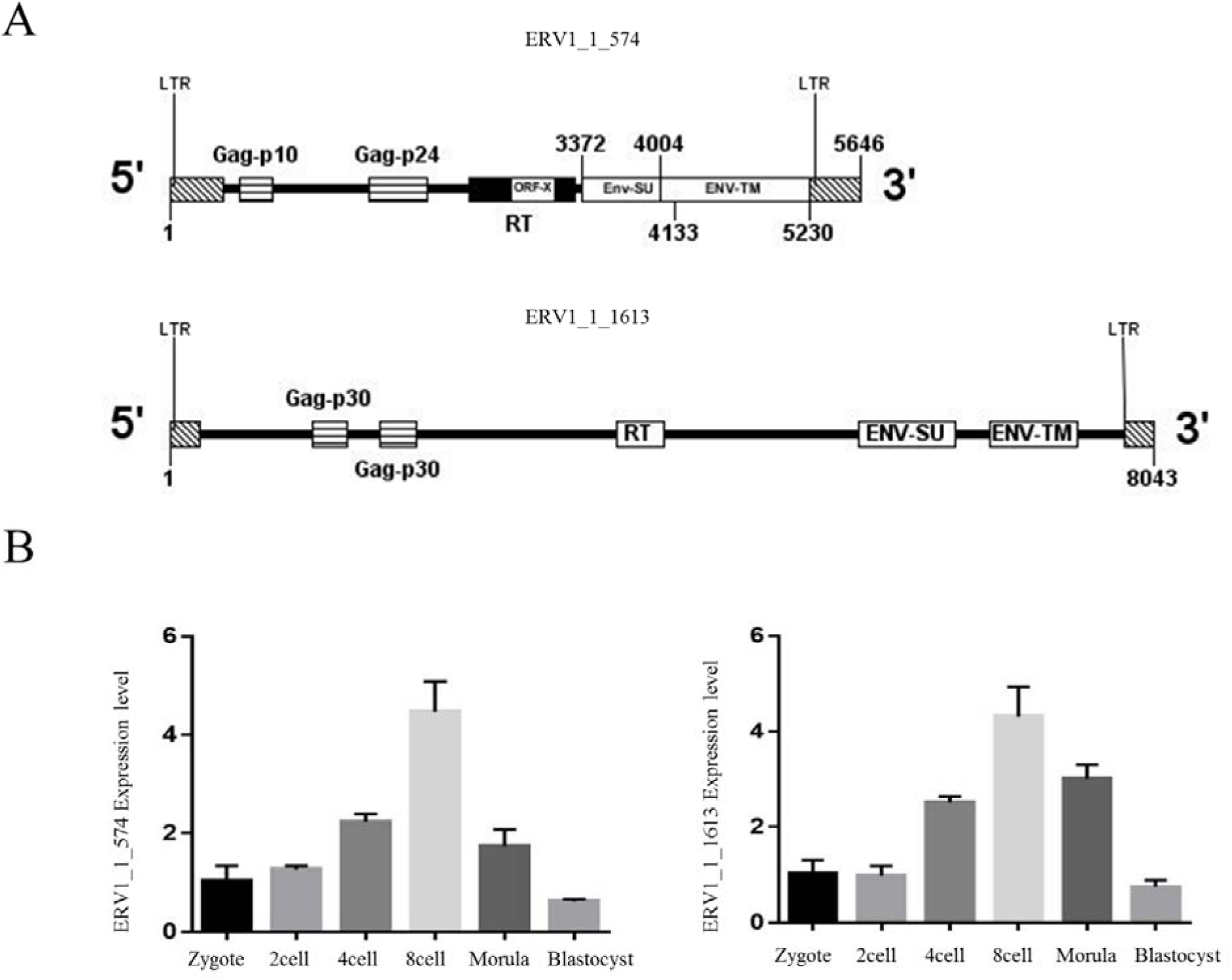
ERV1 has the potential to encode the Gag protein viral capsid. **(A)** ERV1_1_574 and ERV1_1_1613 have Gag and LTR structures at both ends, where Gag_P24 is the core capsid protein. **(B)** Quantitative PCR (qPCR) analysis of ERV1_1_574 and ERV1_1_1613 expression levels in in vitro-fertilized embryos at various stages. Error bars indicate SEM.

TEM results revealed a large number of VLPs distributed in the spaces between blastomeres and in the perivitelline space **(Fig. 4A)**. After ERV1_1_574 knockdown, we found that the number of VLPs was significantly decreased both in the space between blastomeres **(Fig. 4B)** and in the perivitelline space **(Fig. 4C)** in 8-cell stage embryos (*P*< 0.01). These results showed that ERV1_1_574 encoded the Gag protein domain to form VLPs in goat early embryos.

**Figure 4.**
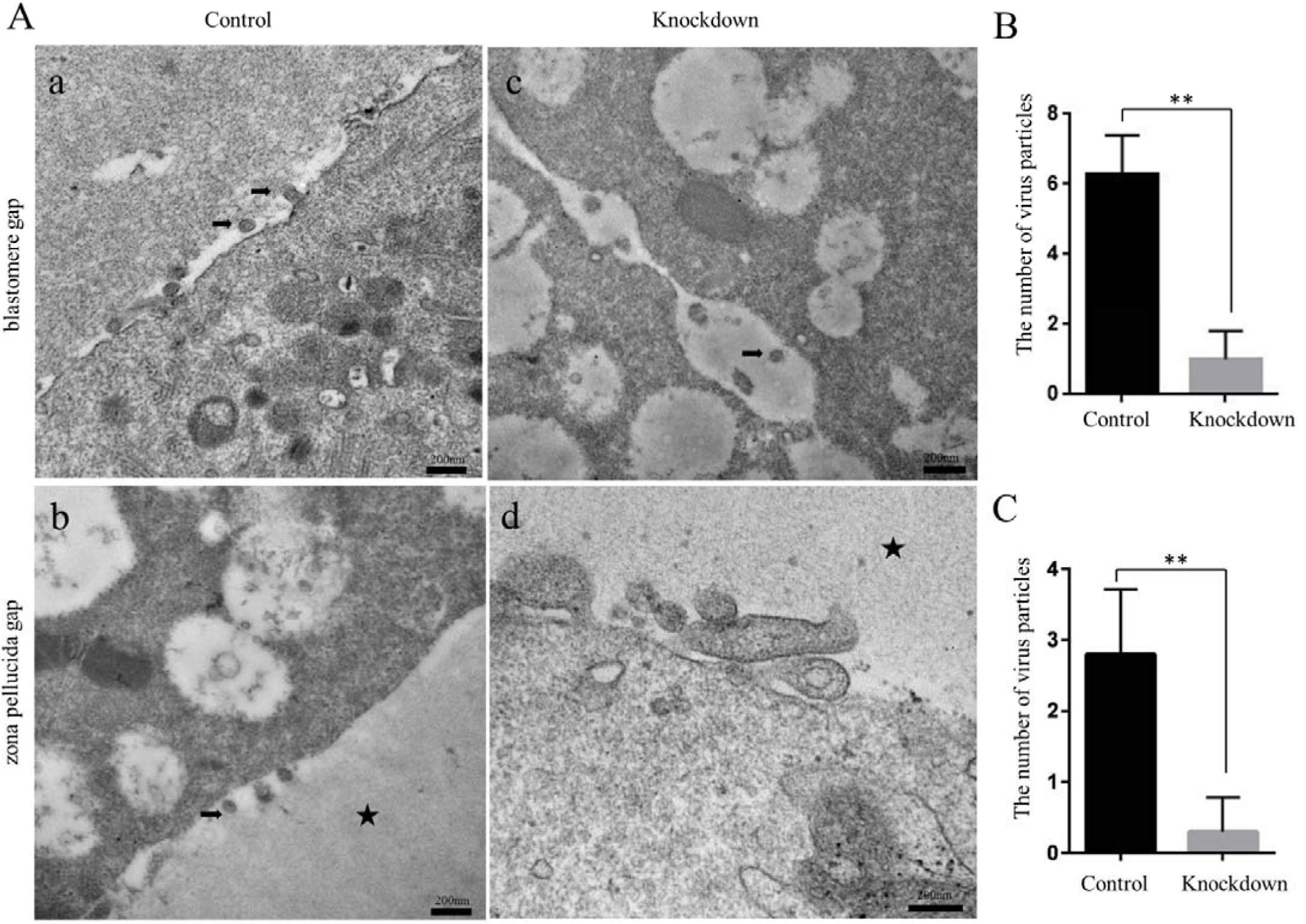
There are a large number of ERV1-derived virus-like particles in early embryos. **(A)** Electron micrographs of Gag particles from the 8-cell stage of goat embryos. Virions appear mainly in the blastomere space and between the blastomere and the zona pellucida. Arrows: Gag particles; stars: zona pellucida. Scale bar: 200 nm. **(B)** The number of Gag particles in the blastomere space. Error bars, SEM. \*\**P*<0.01. **(C)** Number of Gag particles in the gap between blastomere and zona pellucida. Error bars, SEM. \*\**P*<0.01.

### ERV1_1_574 is necessary for early embryo development

We performed RNAi experiments to verify how ERV1 affected early embryonic development **(Fig. 5A)**. The results showed that silencing of ERV1_1_574 significantly reduced the blastocyst development rate, while knockdown of ERV1_1_1613 had no significant effect on embryonic development **(Fig. 5B and C)**. Interestingly, we found that the densification of morula in the ERV1_1_574 knockdown group was significantly affected **(Fig. 5D)**, and few embryos developed beyond sixteen cells to the morula or blastocyst stage **(Fig. 5E and F)**. We performed blastocyst quality assessment experiments through CDX2 staining and found that the total number of blastocyst cells and the number of trophoblast cells in the knockdown group were significantly reduced (*P*<0.05), but the number of inner cell masses was not significantly different **(Fig. 6A and B)**. The apoptotic staining experiments revealed that apoptotic cell numbers in blastocysts were significantly increased in the knock-down group relative to the controls (*P*<0.001) **(Fig. 6C and D)**.

**Figure 5.**
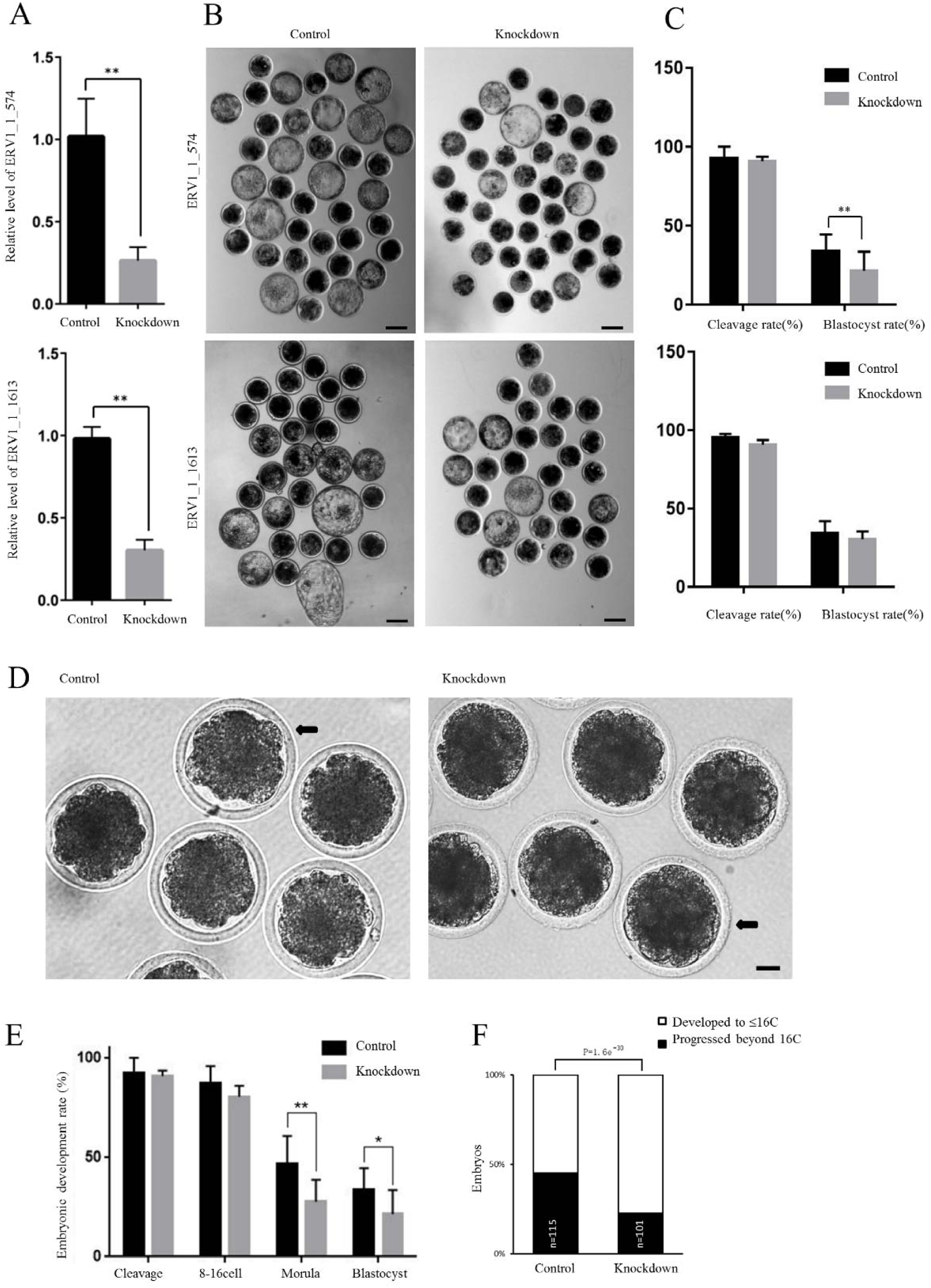
Knockdown of ERV1_1_574 affects early embryonic development. **(A)** Expression of ERV1_1_574 and ERV1_1_1613 at the 8- to 16-cell stage of IVF embryos developed from siRNA/C-injected zygotes. Error bars = SEM. \**P*< 0.05, \*\**P*< 0.01. **(B)** Representative images of control embryos and those siRNA-injected as blastocysts. Scale bar = 100 μm. **(C)** Development rate of the cleavage and blastocysts in siRNA/C-injected embryos. Error bars = SEM. \**P*< 0.05; \*\**P*< 0.01. **(D)** Degree and morphology of morula densification in siRNA/C-injected embryos. Scale bar = 20 μm. **(E)** Development rate of cleavage, 8- to16-cell stage, morula and blastocysts in embryos. Error bars = SEM. \**P*< 0.05; ** *P*< 0.01. **(F)** Developmental embryonic progression after siRNA/C injections. χ^2^ P values were calculated for the developmental rate of embryos injected with siRNA.

**Figure 6.**
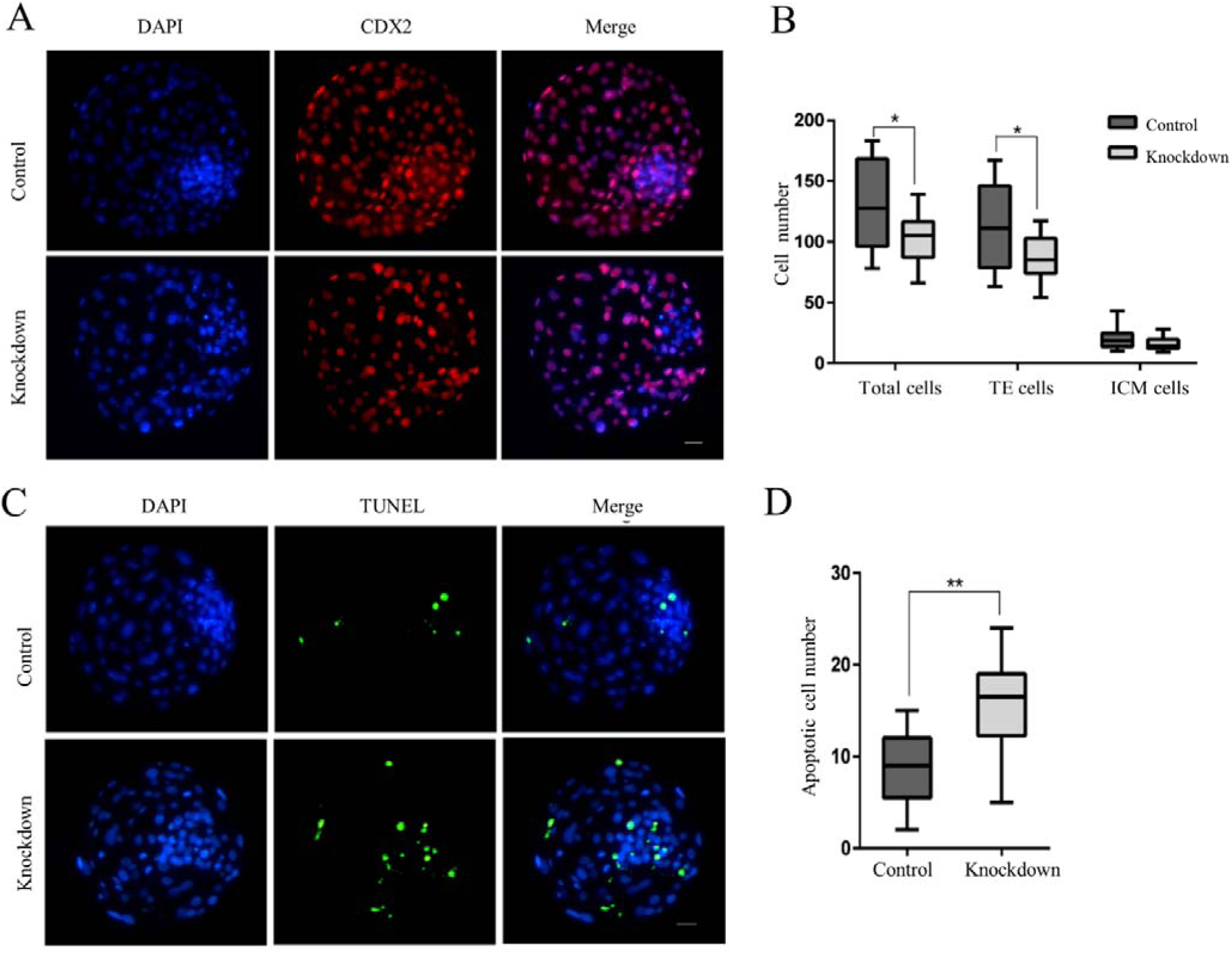
Knockdown of ERV1_1_574 affects blastocyst quality. **(A)** Staining for CDX2 (*red*) and DAPI (*blue*) in blastocysts developing from siRNA/C-injected zygotes. Scale bar = 15 μm. **(B)** Box plots show total cell numbers, TE cell numbers and ICM cell numbers of goat blastocysts from siRNA/C-injected zygotes. \**P*< 0.05 (Student’s *t* test). **(C)** Apoptotic cells (*green*) showing nuclei (*blue*) of blastocysts developing from siRNA/C-injected zygotes. Scale bar = 15 μm. **(D)** Box plots indicate the apoptotic cell number (TUNEL-positive) of goat blastocysts from the siRNA/C-injected zygotes. \*\**P*<0.01 (Student’s *t* test).

### ERV1_1_574 regulated embryo compaction and trophoblast differentiation-related genes

To elucidate the mechanism of ERV1_1_574’s effect on embryo development at the gene level, we performed transcriptome sequencing analysis of morula embryos and obtained 351 differentially expressed transcripts **(Fig.7A)**, of which 188 were upregulated and 163 were downregulated **(Fig. 7B)**. KEGG analysis **(Fig.7C)** and GO enrichment analysis **(Fig.7D)** showed that differential genes were enriched in cell adhesion, cell junction and other pathways. RT-qPCR was used to validate the expression levels of the DEGs, CX43 (a connexin-encoding gene) and CDX2 (embryo lineage differentiation gene), associated with embryo compaction and trophoblast differentiation. Knockdown of ERV1_1_574 significantly decreased the expression of CX43 and CDX2 in morulas **(Fig.7E)**. These results indicated that ERV1_1_574 affected embryonic development by regulating the expression of embryo compaction and trophoblast differentiation-related genes.

**Figure 7.**
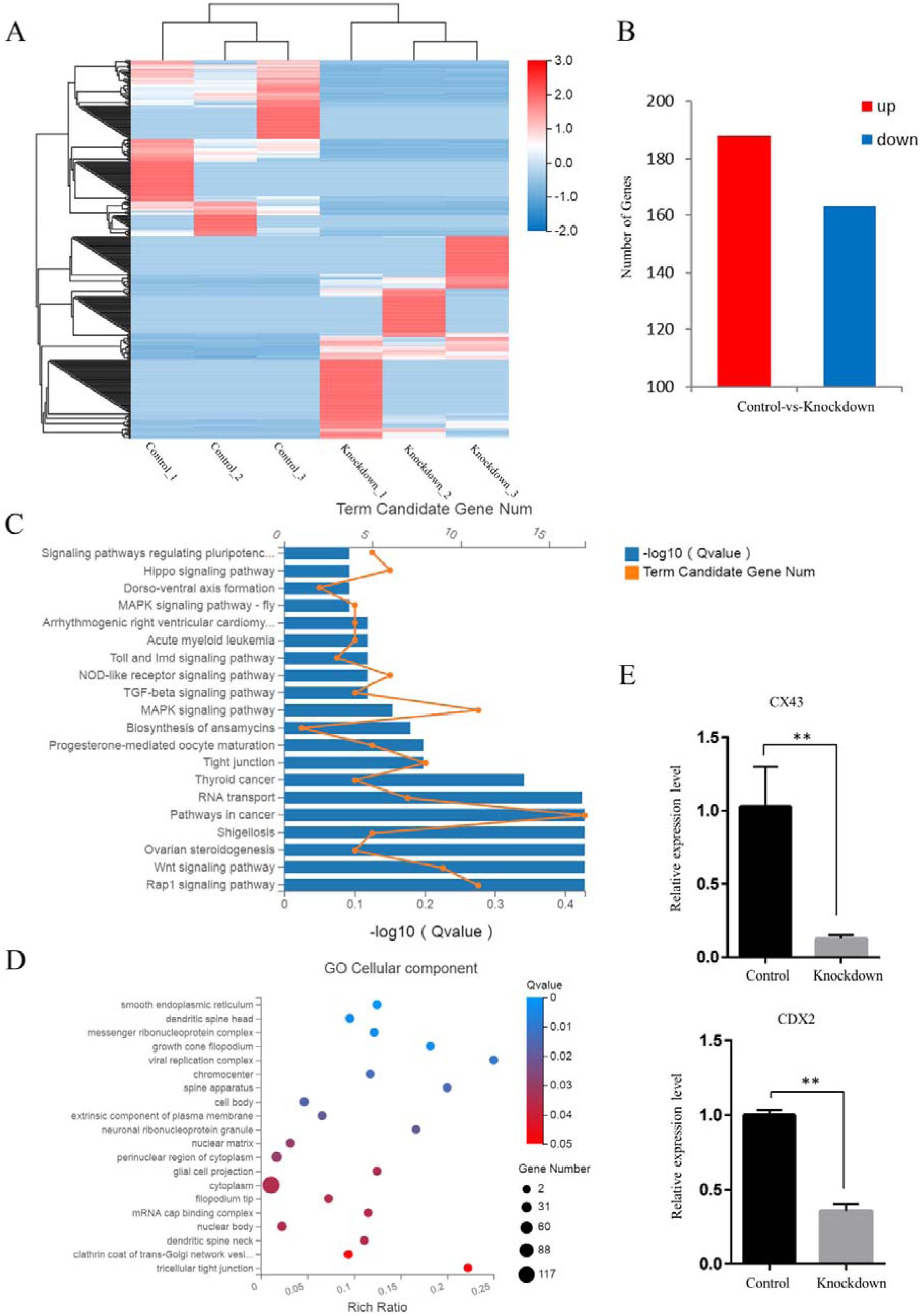
Influence of ERV1_1_574 on embryo densification is related to gap-junction protein. **(A)** Hierarchical clustering analysis of 351 differentially expressed lncRNAs and mRNAs between the control group and the knockdown group at the morula stage. Data are from FPKM. Red: high expression. Blue: low expression. **(B)** Compared with the knockdown group, 188 genes were upregulated and 163 genes were downregulated in the control group. **(C)** KEGG pathway terms are displayed for 351 DEGs in morulae of IVF embryos following siRNA/C microinjections. **(D)** GO analysis of DEGs in morulae of IVF embryos following siRNA/C microinjections. **(E)** Expression of CX43 and CDX2 in morulae of in vitro-fertilized embryos developed from siRNA/C-injected zygotes. Error bars indicate SEM. \*\**P*< 0.01.

## Discussion

ERVs constitute a large number of TE elements in eukaryotic genomes (Mager & Stoye, 2015). Systematic elucidation of the expression and function of ERV elements is useful for revealing the regulatory mechanism of early embryonic development (Gifford *et al*, 2013; Friedli & Trono, 2015; Hutchins & Pei, 2015). In this study, we constructed a TE database and comprehensively analyzed the expression patterns of ERVs during early goat embryo development. We screened ERV1 for its potential to encode a gag protein and detected the presence of VLPs in goat embryos. We found that ERV1_1_574 knockdown caused the arrest of embryos mostly at the 8- to16-cell stage; therefore, we explored the possible biological function of ERV1_1_574 on early embryonic development in goats.

The subfamily and function of reactivated ERV elements in early embryos were different among mammal species. In mouse, MuERV-L was an important marker of zygotic genome activation with important functions in early embryo development and establishment of pluripotency (Kigami *et al*, 2003; Macfarlan *et al*, 2012). In early human embryos, HERVH was mainly expressed in the inner cell mass and promoted embryonic stem cells through the encoded lncRNA, HPAT5 (Durruthy-Durruthy *et al*, 2016; Wang *et al*, 2016). ERV1s were highly expressed in early embryos, but their function and regulation were unclear in bovines (Bui *et al*, 2009). Our results showed that ERV1_1_574 was highly expressed specifically during EGA, and that ERV1_1_574 knockdown significantly reduced embryo compaction and trophoblast differentiation in goats.

ERV elements were involved in regulating gene expression during embryonic development through various pathways, including promoter/enhancer activity, non-coding RNA (lncRNA), functional proteins, and epigenetic modification (Chuong *et al*, 2017; Hendrickson *et al*, 2017). In previous studies, HERV-K LTR was shown to contain multiple transcriptional start sites (TSS), and alternate TSS were part of the active regulation of gene transcription directed by LTRs (Fuchs *et al*, 2011; Persson *et al*, 2016). The ERV-derived lncRNA, LincGET, influenced inner cell mass (ICM) development and induced cell fate decisions (Wang *et al*, 2018). The HERV-W env protein acted as a fusion protein to promote the formation of syncytiotrophoblast cells (Frendo *et al*, 2003). HERV-K, which exists in repressed chromatin regions, has a strong association with H3K9me3 and is able to both activate and suppress gene expression (Campos-Sánchez *et al*, 2016). Our study found that ERV1_1_574 affected the development of early embryos by regulating the expression of CX43 and CDX2, which are genes related to embryo compaction and trophoblast differentiation. We also determined whether ERV-encoded Gag protein could regulate embryonic development.

In recent years, the functions and regulatory mechanisms of various ERV-derived non-coding RNAs have been unraveled (Gerdes *et al*, 2016); however, less is known about the function and regulation of Gag proteins encoded by complete ERV sequences. Previous studies revealed that the complete ERV element, MuERV-L, could produce Gag protein and form VLPs in 2-cell stage mouse embryos (Macfarlan *et al*, 2012). In addition, the envelope protein of endogenous Jaagsiekte sheep retrovirus (enJSRVs) was able to regulate trophectoderm growth and differentiation in peri-implantation embryos (Dunlap *et al*, 2006). Here, we confirmed the presence of Gag protein VLPs in early goat embryos, and established that ERV1_1_574, had the ability to express Gag protein, and significantly affect the rate of embryo development, and the rate and quality of blastocyst formation. Whether ERV1_1_574 regulated this process alone, or through Gag proteins, is still under investigation.

The Gag proteins produced by ERVs have specific biological functions in placental development. The Fv1 (Friend virus susceptibility-1) gene, encoded by the mouse MuERV-L Gag protein gene, can limit infection by the mouse leukemia virus MuLVs (Sanz-Ramos & Stoye, 2013). In sheep, the Gag protein encoded by enJSRV restricted the replication of exogenous viruses by blocking their interaction with receptors (Arnaud *et al*, 2007). Recent studies have shown that VLPs derived from ERVs can enclose their own RNAs and transmit them between cells, suggesting that virus particles derived from ERVs are capable of mediating cell communication. During Drosophila oogenesis, transposons were passed from supporting nurse cells to eggs by means of microtubules to promote oocyte development, proving that transposons can be passed from cell to cell (Wang *et al*, 2018). In neurons, the Arc gene encodes a Gag-like protein that forms VLPs, and the Arc gene mRNA is delivered to other neuronal cells in the form of exosomes (Pastuzyn *et al*, 2018). Additional studies will need to be done to determine whether the Gag protein encoded by ERV1 can be assembled into VLPs that can act like exosomes to encapsulate specific RNAs and pass through the cytomembrane in early goat embryos to mediate communication between blastomeres.

In summary, our study demonstrated that ERV1 had the potential to encode the Gag protein domain to form VLPs that regulate trophoblast cell differentiation in early goat embryos. Although only a subset of ERVs has been studied during early embryonic development, our findings highlight the unexplored functions of ERVs in regulating embryonic development.

## Materials and Methods

### Establishment of goat TE database

According to the latest goat genome data, LAMP (Linux Ubuntu Server 12.04, Apache 2, MySQL Server 5.5, Perl 5.16.3/ PHP 5.3) was used to construct a database of TEs from the goat genome. The TE data were stored as MySQL tables. Common gateway interface (CGI) programs were adapted using Perl, JavaScript and PHP programming languages. The JBrowse genome browser is an embedded genome browser produced with HTML5 and JavaScript that was employed to manipulate and depict the positional relationships between genes and TEs in the goat database (Skinner et al., 2009). The establishment of the goat TE database was mainly based on the signature, homology, and Denove methods. Predict LTRs (long terminal repeats), Helitron MITEs (miniature inverted repeat transposable elements), LINEs (long interspersed nuclear elements), SINEs (short interspersed nuclear elements), and TIRs (terminal inverted repeats) were identified. The filter sequence was redundant, and the filter criterion was identity >90%. LTRs and LINEs were used for superfamily identification by direct comparison with the Repbase database, and family identification was based on the 80-80-80 principle for classification. The coverage and classification of transposon elements in the goat genome was performed using RepeatMasker (http://www.repeatmasker.org, v 4.0.3). The Repbase update collection was down-loaded from http://www.girinst. org/repbase/index.html (Jurka et al., 2005). The position of the transposon sequence in the genome was obtained by using the visualization software from UCSC. The Jukes-Cantor step-length was calculated and the appearance of the evolutionary tree was further edited by Mega v7 (Kumar et al., 2016). Lastly, the potential active elements in the open reading frame were identified via CLC sequence Viewer 5 (CLC Bio). The data used in this study was down-loaded from the goat genome database https://www.ncbi.nlm.nih.gov/genome/?term=goat.

### Goat embryo collection and culture

The goat ovaries were collected from an abattoir, placed in saline containing 100 U/ml penicillin/ streptomycin at 20℃, and transported to the lab within six hours. The organs were washed 3× and incubated in TCM-199 (Gibco-BRL, NY, US) with 25 mM HEPES, 10% fetal bovine serum, and 2 IU/mL heparin. Follicles (2-6 mm diameter) were removed, and cumulus-cell oocyte complexes (COCs) with intact/dense cumulus cells were removed under a stereo-microscope. After washing with phosphate-buffered saline, 100 COCs were pipetted into four-well plates and cultured in maturation medium (TCM-199 supplemented with 10% FBS, 0.2 mM sodium pyruvate, 0.075 IU/mL human menopausal gonadotropin, 1 µg/mL 17β-estradiol, 10 ng/mL epidermal GF, and 1% insulin/transferrin-selenium) for about 24 hours at 38°C in a CO_2_ incubator.

Commercial cryopreserved semen (SKXing Breeding Biotechnology, Inner Mongolia, China) was employed for fertilization. Fifty microliter sperm aliquots (2 × 10^6^ sperm/mL) were incubated with 40 to 50 COCs in 400 µL BO-IVF medium (IVF Bioscience, Falmouth, UK). After twelve hours, the COCs were removed and the zygotes transferred into 400 µL BO-IVC medium (IVF Bioscience, Falmouth, UK) and layered with mineral oil. Embryos were removed at specific stages for experiments.

### RNA-seq

The single-cell RNA-seq technology (SUPeR-seq) developed by Tang et al. was performed to profile the whole transcriptome (Fan et al., 2015). Briefly, the three best quality embryos at each developmental stage were disrupted in single-cell sequencing lysis buffer (blastocysts were not used to construct the sequencing library for technical reasons). RNA was reverse transcribed into cDNA using the T15N6 primers from the SuperScript III kit. The amplified and purified cDNA product was sheared into 150-350 bp fragments by Covaris S2. Fragmented DNA was amplified using the TruSeq DNA library building kit. All libraries were sequenced on an Illumina platform to generate the raw data. Approximately 11.4 million reads were obtained per sample.

The sequencing data were filtered with SOAPnuke (v1.5.2) (Li et al., 2008), and clean reads were mapped to the goat assembly ARS1 genome using HISAT2 (v2.0.4) (Kim et al., 2015). Transcripts were assembled by StringTie (Pertea et al., 2015) to generate novel, known transcripts. Fragments per gene were counted using HTSeq-count (version 0.12.4) (PMID: 25260700) from aligned reads. Fragments per kb of exon model per million mapped fragments (FPKM) were determined using DESeq2 (v1.4.5) (Love et al., 2014) after data normalization. To compare expression of genes between different developmental stages, normalization of reads coverage and differential gene expression analysis at the different developmental stages were performed using DEseq2. The threshold of differential expressed genes was *P*< 0.05, |fold change| > 1.

### Analysis of the source transcripts of TEs

Protein-based RepeatMasking in Repeatmasker was used to compare the transcriptome sequence with all transposon element libraries and obtain dynamic expression levels of transcripts related to transposon elements.

### Predictive analysis of ERV1 domains

GenomeTools 1.5.7 was used to analyze the structure of ERV1 and to predict the structure of ERV1 with domains >3.

### RT-qPCR

Embryos at the zygotic, two-, four-, and eight-cell stages, and morulae were harvested, respectively, at 1, 2, 3, and 5 days after fertilization or activation on day 0. The PrimeScript^TM^ RT reagent kit with gDNA eraser (Takara) was used to convert total RNA to cDNA. SYBR^®^ Premix Ex Taq^TM^ II and a one-step real-time PCR system (Applied Biosystems) was used to do reverse-transcription quantitative PCR (RT-qPCR). The primer sequences are shown in Table 3, and data are compared as fold-change = 2^−ΔΔCt^ means ± SD.

### Embryonic RNA interference (RNAi)

The siRNA against the sheep ERV1_1_574 transcript was created and synthesized by GenePharma (Shanghai, China) (Table 4). The siRNA was aliquotted at 20 µM and kept at [80°C. About 10 pL of siRNA was injected into a presumed fertilized egg (collected 20 hours after IVF). The siRNA was microinjected under a microscope (Axio Observer D1, Zeiss) equipped with a micro-manipulation device (TransferMan NK2, Eppendorf) and a micro-injector (FemtoJet, Eppendorf). A volume of 1μL of siRNA was drawn from the bottom of the tube through a micro-sampler (Eppendorf). During microinjection, the embryo was inside a 100 μL droplet of M199/HEPES (Gibco) plus 10% fetal bovine serum.

**Table 4.**
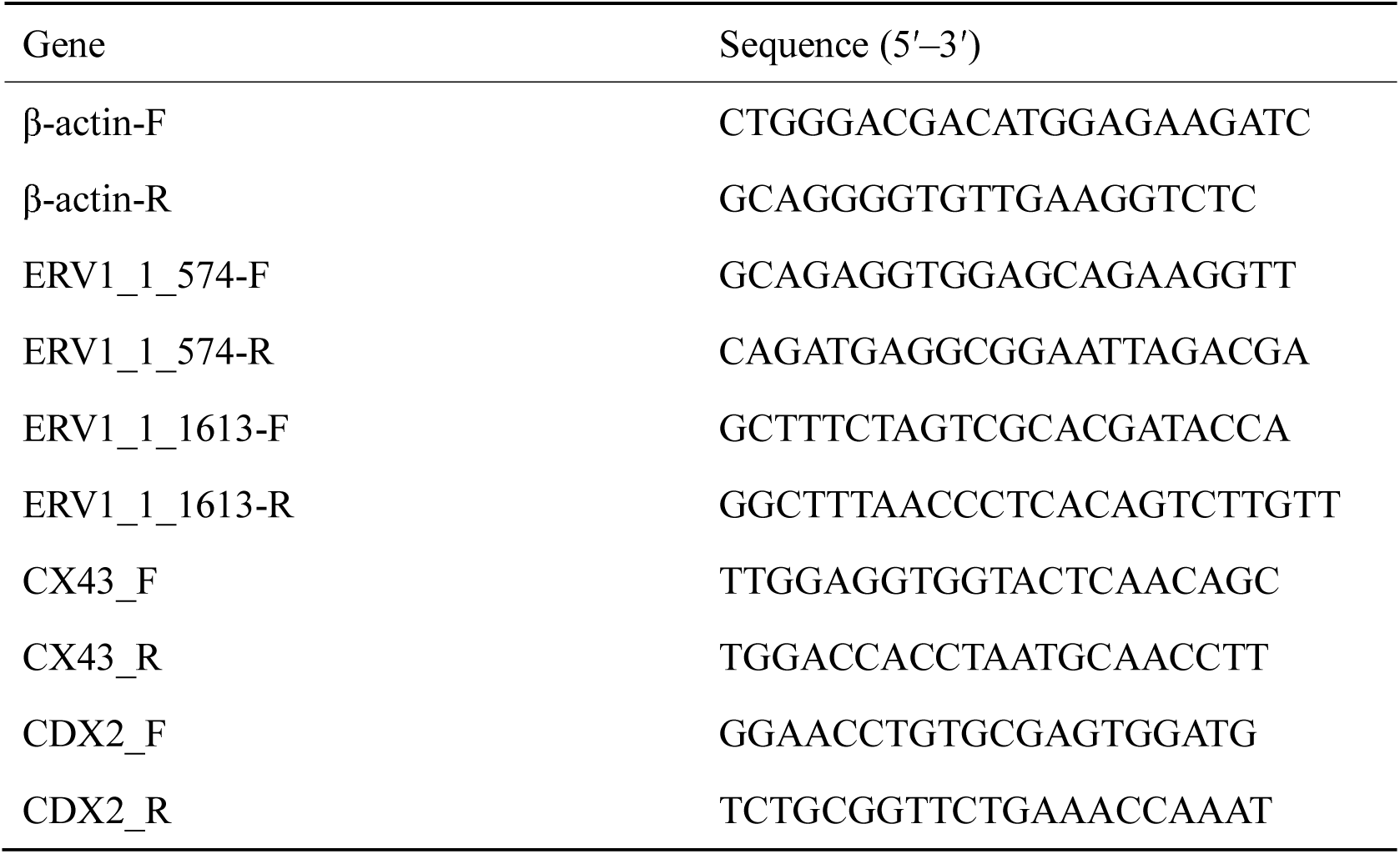
siRNA sequences for RNA interference.

**Table 5.**
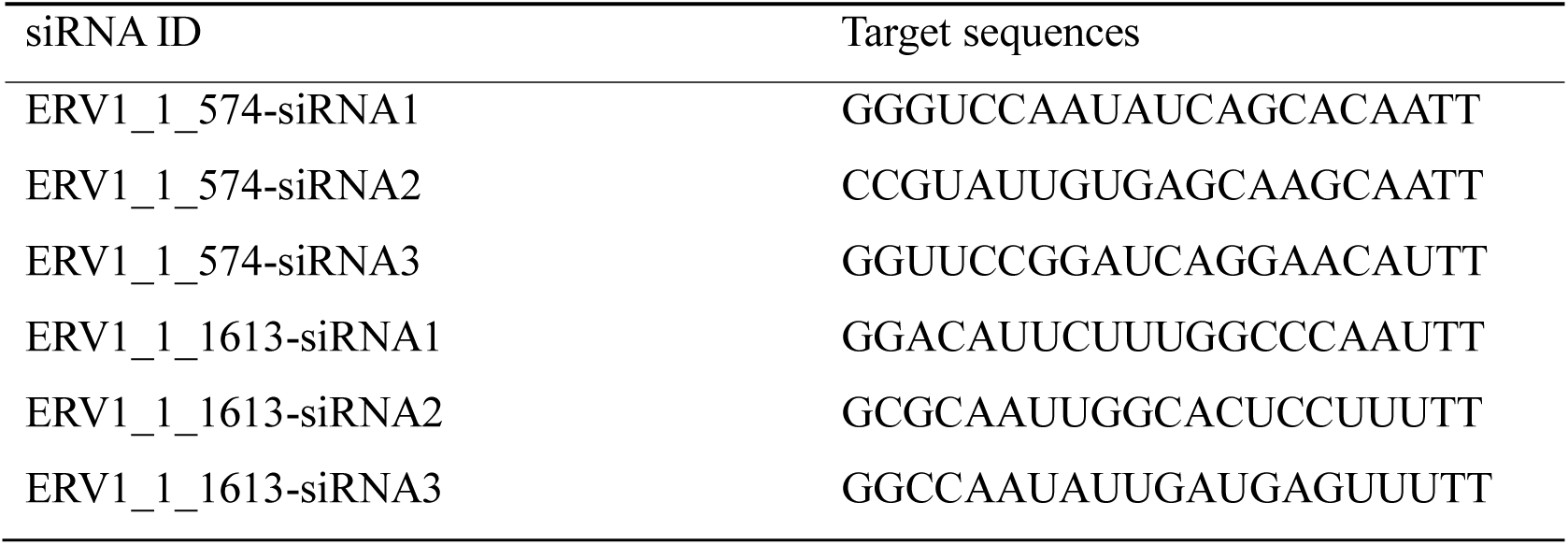
Primer sequences for RT-qPCR.

### Differential staining and apoptosis analysis of early embryonic cells

Experimentally treated blastocysts and control blastocysts were collected on day 7, fixed with 4% paraformaldehyde at room temperature for 4 hours, and permeabilized with 0.1% Triton X-100 at room temperature for 20 minutes. The blastocysts were blocked in immunostaining blocking medium at 4°C overnight followed by incubation with anti-CDX2 (BioGenex Inc., US) at 4°C overnight. After washing, the blastocysts were incubated with AlexaFluor 555-conjugated donkey anti-mouse IgG for 2 h in the dark at room temperature. After washing again, the nuclei in the samples were stained with DAPI (Beyotime, China) and the stained cells were imaged under a fluorescence microscope. The total number of blastocyst cells, trophoblast cells and inner cell clusters were counted.

Another group of blastocysts was collected for measuring apoptosis. After fixing and permeabilization, the blastocysts were transferred to a solution of 45 µL of E-buffer, 5 µL dNTP Nad mix, and 1µL rTdT, and incubated at 37°C in the dark for 1 hour. Reactions were terminated by addition of 2X SSC, nuclei were stained with DAPI, and apoptosis was observed under a fluorescence microscope. The total number of blastocyst cells (DAPI) and apoptotic cells (FITC) were counted and the apoptosis rate was determined. The TUNEL assay kit used in the experiment was from Promega Corporation.

### Transmission electron microscopy

Eight-cell stage embryos were fixed with 2.5% glutaraldehyde for 1 hour at 4°C. The agar embedding method was used to embed 30-40 embryos in a 2 mm x 2 mm x 2 mm agar block. The agar blocks were rinsed three times with 0.1 M phosphate buffer, fixed with 1% osmic acid solution for 2-4 h, then dehydrated twice in graded ethanol solutions: 30%, 50%, 70%, 80%, 90%, and 100%. Samples in 100% ethanol were immersed in LR-White embedding agent (3:1) for 2 h, 1:1 for 8h, 1:3 for 12 h, and lastly, 100% LR-White for 24 h, overnight. The samples were placed in the embedding polymerizer for automatic polymerization and then sectioned, first as semi-thin sections and, after positioning, as ultra-thin sections. The ultra-thin sections were floated onto copper meshes, stained with uranyl acetate, then with lead citrate, dried naturally and observed under a transmission electron microscope.

### Analysis of ERV1 expression interference by embryonic gene expression

Heatmaps of the various samples were constructed using P-heatmap (v1.0.8) based on gene-expression profiles. Differential expression was determined using DESeq2 (v1.4.5) (Fan et al., 2015) with *q* value ≤ 0.05. For additional insights into the changes of phenotypes, GO (http://www.geneontology.org/) and KEGG (https://www.kegg.jp/) enrichment analysis of annotated DEGs was done using the P-hyper package (https://en.wikipedia.org/wiki/Hypergeometric_distribution) based on the hypergeometric test. The significance levels of terms and pathways were corrected according to q values with a strict threshold of *q*≤ 0.05, using the Bonferroni test.

### Statistical analysis

Appropriate statistics tests were performed using SPSS 22.0 (SPSS Inc., US). Experiments were repeated at least three times, and data are means ± standard error of the mean (SEM). For RT-qPCR, the value of 2^-△△Ct^ was measured to compare the relative gene expression of experimental with control groups. Student’s *t* test or Chi-squared test was used for paired comparison, and one-way analysis of variance was done for multiple comparisons. *P*< 0.05 indicated significance

### Compliance and ethics

All animal procedures and experiments were conducted in accordance with the Guide for the Care and Use of Laboratory Animals (Ministry of Science and Technology of China, 2006), and were approved by the animal ethics committee of Northwest A&F University.

## Acknowledgments

We thank Ronghua Zhang for providing the goat ovaries.

## Author contributions

Wenjing Li: Data curation, Methodology, Formal analysis, Writing. Shujuan Liu: Methodology, Formal analysis. Jianglin Zhao: Methodology, Formal analysis. Ruizhi Deng: Methodology, Formal analysis. Yayi Liu: Methodology, Resources. Huijia Li: Methodology, Resources. Hongwei Ma: Methodology, Resources. Yanzhi Chen: Writing. Jingcheng Zhang: Methodology, Resources. Yongsheng Wang: Methodology, Resources. Jianmin Su: Methodology, Resources. Fusheng Quan: Methodology, Resources. Yong Zhang: Project administration, Xu liu and Yan Luo: Conceptualization. Jun Liu: Conceptualization, Writing, Funding acquisition.

## Supplementary Data Legends

**Figure S1.**
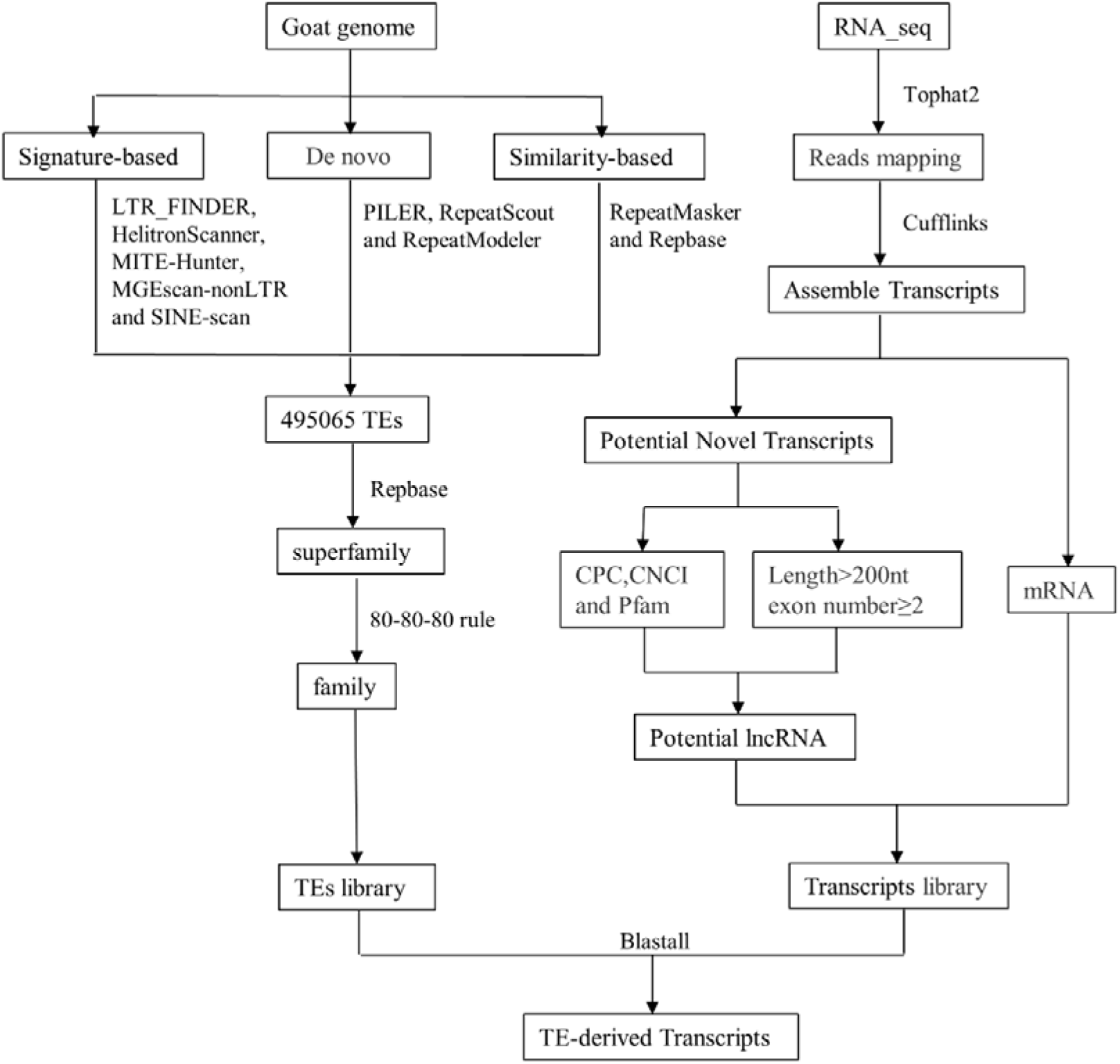
Flow chart of TE-derived transcripts. Left: flow chart of complete TEs. A total of 495,065 complete TEs were identified based on three methods (*ab initio* prediction, structure prediction and homology prediction strategy). All TEs were classified into 21 superfamilies by comparing the custom repeat library with the Repbase. According to the 80-80-80 rule, all putative goat TEs were classified into 926 families. Right: lncRNA and mRNA discovery for RNA-Seq. Bottom, TE-derived transcripts. We used Blastall to blast lncRNA and mRNA with complete TEs.

**Figure S2.**
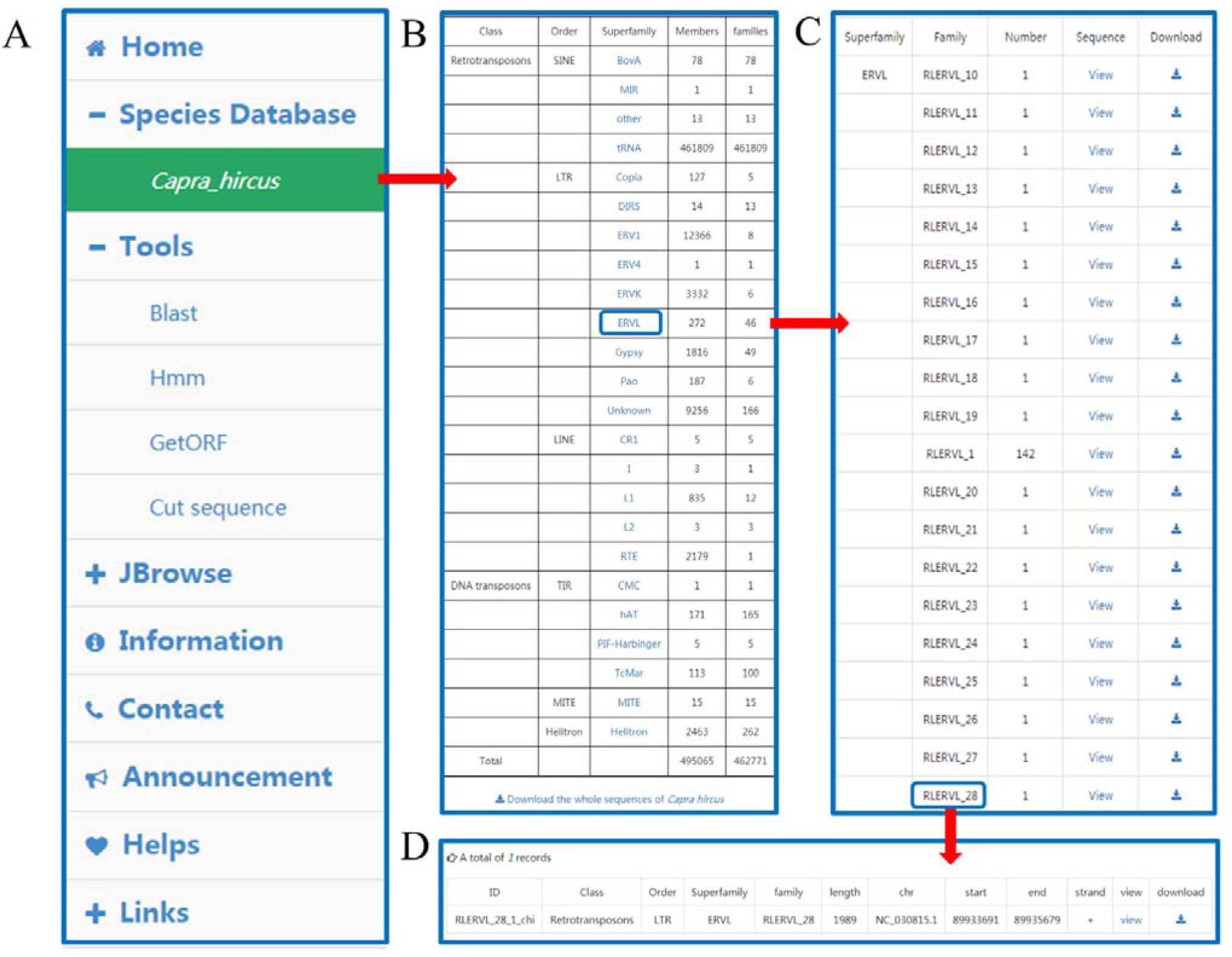
User interface for browsing in GOAT-TEdb. **(A)** Menu of GOAT-TEdb. **(B-D)** Browsing interface of database GOAT-TEdb.

**Figure S3.**
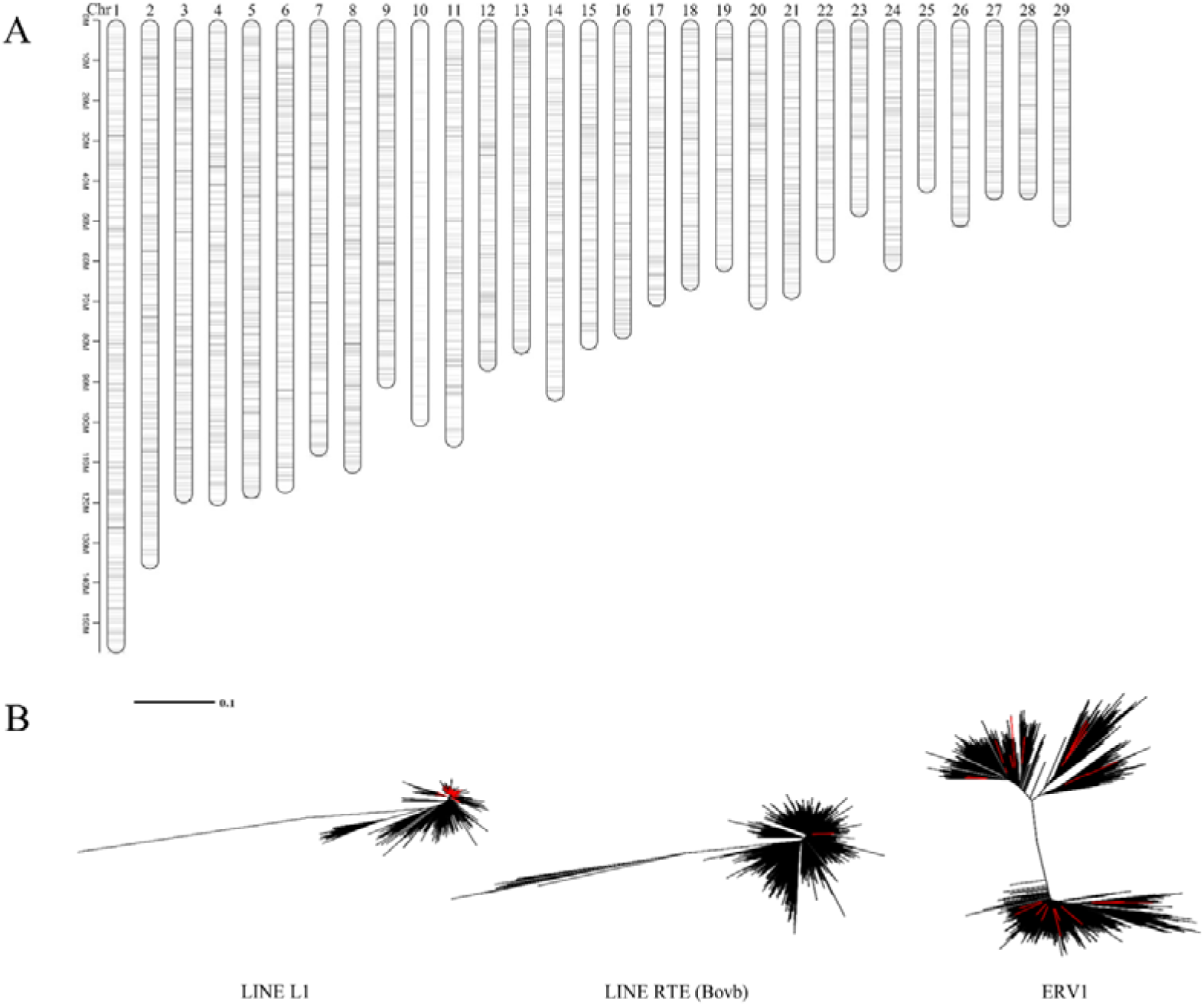
Chromosomal distribution and maximum likelihood trees of TEs. **(A)** Location of goat TEs in the genome. The x axis corresponds to the chromosomes, the y axis to nucleotide coordinates in million base-pairs (Mbp) in the goat genome. **(B)** Intact RTE, L1 and ERV trees. Maximum likelihood trees derived from global alignments of all intact/full-length LINEs and ERV sequences. Red lines depict potentially active LINEs or ERVs based on their ORF content.

**Figure S4.**
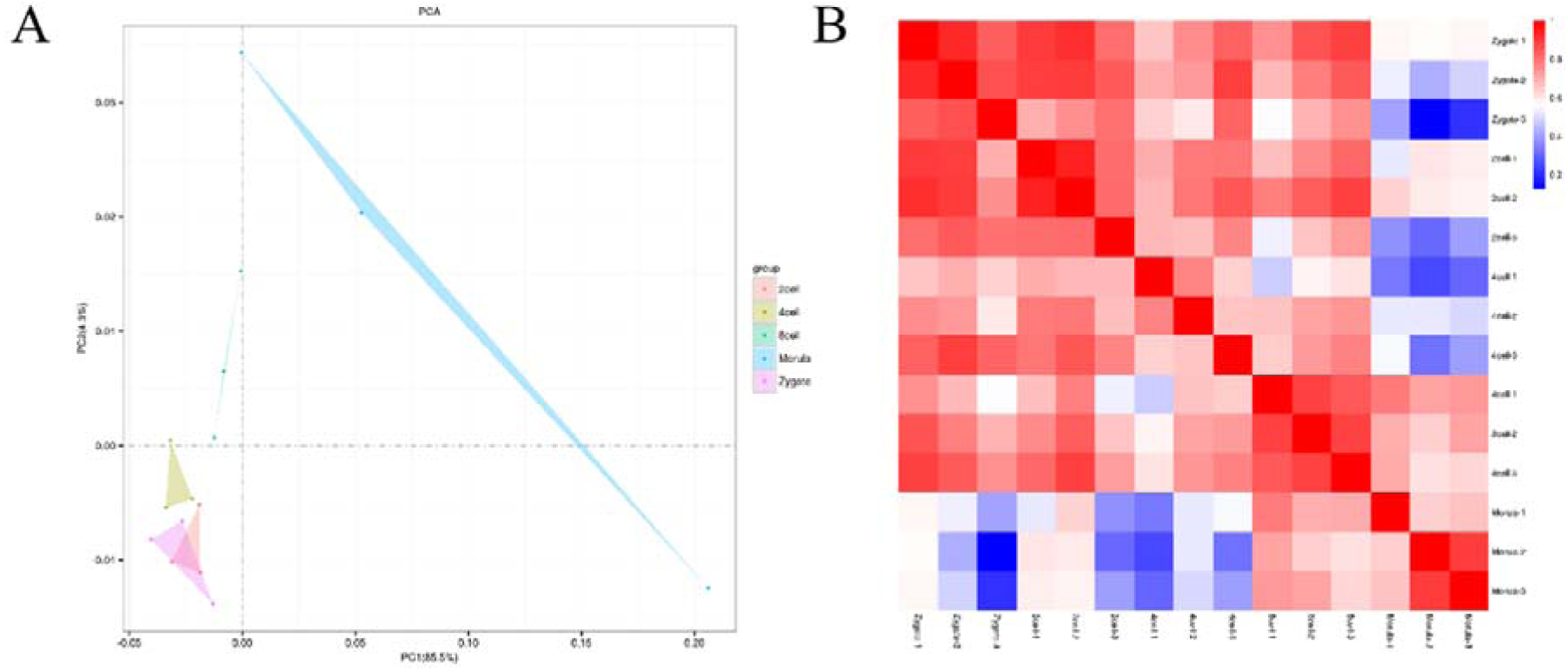
Sample correlation analysis. **(A)** Correlation of single-cell goat sequencing samples. Principal-component analysis of the transcriptomes of single embryos during preimplantation development. **(B)** Pearson correlation-coefficient heatmap of single embryo transcriptomes during preimplantation development.

**Figure S5.**
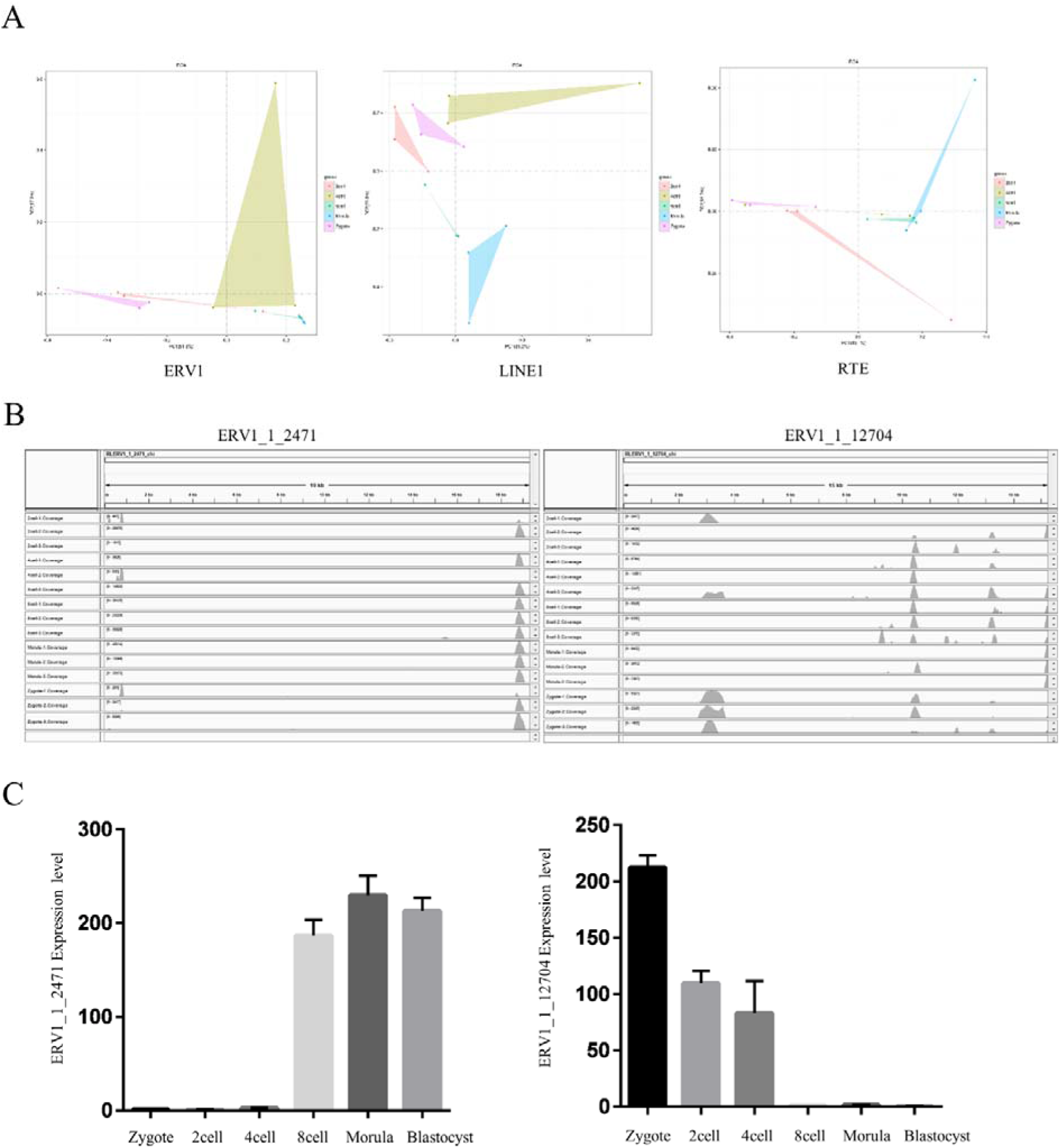
Stage-specific TE expression during early goat embryonic development. **(A)** PCA of ERV1, LINE1 and RTE expression estimates in pre-implantation goat embryos. The TEs showing the largest variation between the stages were chosen. ERV1, LINE1and RTE expression are representative of the different development stages. (B) Normalized RNA-seq for two loci showing stage-specific ERV1 expression (ERV1_1_2471 and ERV1_1_12704). (C) Expression profile of ERV1_1_2471 and ERV1_1_12704. β-actin was used as a control. Error bars = SEM.

**Figure S6.**
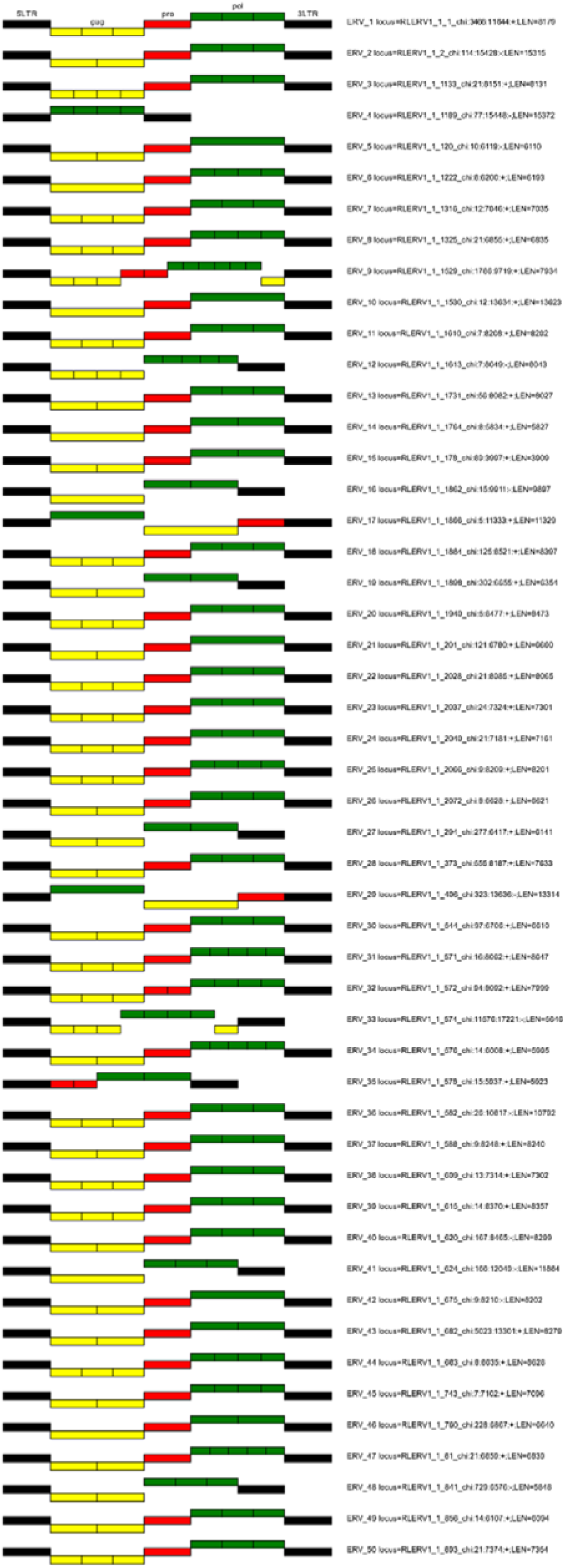
Structural features of ERV sequences. Gag, Pol and Pro indicate ERV repeat regions.

